# The olfactory organ is a site for neuroendocrine modulation of reproduction in zebrafish

**DOI:** 10.1101/2025.07.07.663467

**Authors:** Eugene M. Tine, Ingrid Pinto-Borguero, Constanza Aguirre-Campos, P. Marcela Escobar, Ricardo Fuentes, John Ewer, Kathleen E. Whitlock

## Abstract

Gonadotropin-releasing hormone (GnRH) is one of the most fascinating neuroendocrine peptides: It is essential for maintaining the reproductive state of vertebrates and also displays high sequence homology to the tridecapeptide mating pheromone of the yeast, *S. cerevisiae*. In spite of its highly conserved role in vertebrate reproduction, recent studies in zebrafish show that the loss of function of genes encoding Gnrh isoforms does not cause infertility. Here we first investigated whether Phoenixin, a novel peptide acting in the reproductive pathway of vertebrates, is the hormone that has replaced Gnrh in zebrafish. While loss of function of the *phoenixin* gene affected female differentiation, we observed no defects in fertility. We next reconsidered the GnRH pathway and turned to the natural world, where fishes use waterborne hormones to control reproduction. Thus, we investigated whether exogenous Gnrh affects the hypothalamic-pituitary axis. Here we show that fish isolated from their conspecifics and kept in artificial water devoid of fish odors, mounted sex–appropriate pituitary responses when Gnrh was added to the water. Furthermore, blocking the access of water to the olfactory organs eliminated these responses. We then analyzed Gnrh signaling by knocking out the gene encoding *gnrh-receptor3* (*gnrh-r3*) and, surprisingly, found that fish homozygous for a *gnrh-r3* null mutation were almost completely infertile: males did not produce sperm and females produced but a few mature oocytes. Finally, we found that Gnrh was present in nanomolar concentrations in the water that houses the fish, thus supporting the hypothesis that waterborne Gnrh from conspecifics plays a key role in regulating zebrafish reproduction in the absence of the endogenous ligand.

## Introduction

In vertebrates, the neuroendocrine regulation of reproduction is controlled by gonadotropin-releasing hormone (GnRH), a decapeptide released from neurons in the hypothalamus that triggers the release of follicle stimulating hormone (FSH) and luteinizing hormone (LH) from the anterior pituitary (Gore 2002). In most vertebrates, the Gnrh1 isoform, which is secreted by neurons located in the pre-optic area, is the principal neuroendocrine peptide controlling the release of gonadotropins. Teleost fishes usually have three isoforms of Gnrh, which are encoded by separate genes: *gnrh1*, *gnrh2* and *gnrh3*, where Gnrh3 is the fish-specific isoform (Sherwood, Eiden et al. 1983; Powell, Zohar et al. 1994, Fernald and White 1999, Okubo and Nagahama 2008, Tostivint 2011, Sukhan, Kitano et al. 2013). The neurons expressing these peptides are found in different locations in the brain and have varying functions (White and Fernald 1998; Lethimonier, Madigou et al. 2004), where the hypophysiotropic Gnrh neurons either contain Gnrh1, or Gnrh3, or both (Fujimori, Sugimoto et al. 2024).

### Loss of function of gnrh does not result in infertility

The genome of zebrafish (*Danio rerio*) lacks the gene encoding Gnrh1 (Whitlock, Postlethwait et al. 2019), suggesting that in this species the regulation of the reproductive axis might be under the control of Gnrh3 (Karigo and Oka, 2013). However, disabling the *gnrh2* and *gnrh3* genes (Spicer, Wong et al. 2016; Marvel, Spicer et al. 2018) as well as *kiss1* and *kiss2*, two upstream regulators of Gnrh secretion (Liu, Tang et al. 2017), does not affect the fertility of zebrafish. Thus loss of *gnrh* function either through evolutionary gene loss (*gnrh1*; (Whitlock, Postlethwait et al. 2019) or gene editing (*gnrh2*, *gnrh3*: (Spicer, Wong et al. 2016; Marvel, Spicer et al. 2018), does not affect the fertility or the fecundity of adult zebrafish. Although qPCR analysis of reproductive and hypophysiotropic peptides in *gnrh3/gnrh2* double knockout fish revealed that compensatory changes in gene expression may have occurred (Marvel, Spicer et al. 2018), to date only the gene encoding secretory granule protein secretogranin-2 (Scg2), which is a precursor for the neuropeptide secretoneurin (SN), appears to play role in zebrafish reproduction (Ahamed, Hassan et al. 2024; Peng, Lu et al. 2025). Indeed, animals mutant for *scg2* showed abnormalities in male reproductive behavior, resulting in a marked decrease in the number of eggs laid (Mitchell, Zhang et al. 2020). Interestingly, mutations in *scg2a* had no significant impact on the performance of mating behavior in Medaka (Fleming, Tachizawa et al. 2023). Thus, to date none of the genes that could compensate for the loss of *gnrh3* and *gnrh2* have been shown to be essential for fertility.

## Results

### Have zebrafish co-opted a different peptide into the reproductive pathway?

Phoenixin (Pnx) (also known as Small Integral Membrane Protein 20, *Smim20*), is a preprohormone with five predicted cleavage products where the amidated 14 amino-acid (Pnx-14) and the 20 amino-acid (Pnx-20) products are biologically active (Yosten, Lyu et al. 2013; Palasz, Janas-Kozik et al. 2018; Suszka-Switek, Palasz et al. 2019). This highly conserved peptide hormone plays a role in regulating the hypothalamic-pituitary axis, (Yosten, Lyu et al. 2013; Palasz, Janas-Kozik et al. 2018; Suszka-Switek, Palasz et al. 2019), and also has metabolic functions (Muzammil, Barathan et al. 2024). In addition, it has been proposed to play a role in zebrafish reproduction (Whitlock, Postlethwait et al. 2019; Rajeswari and Unniappan 2020; Ceriani, Calfún et al. 2021) by replacing the function of Gnrh1 in the brain-pituitary-gonad axis. To determine the physiological role of zebrafish phoenixin (zPnx-20) in the expression of pituitary gonadotropes mRNAs (*lhb* and *fshb*), adult animals were challenged by injection of zPnx-20 peptide. The resulting increases in *lhb* mRNA and decreases in *fshb* mRNA levels in the pituitary of males (**Supp. Fig. 1**) are consistent with previously published results showing that intraperitoneal injection of Phoenixin induces an increase in *lhb-receptor* and a decrease in *fshb-receptor* expression in the testis of male zebrafish. In contrast, results showing increases in *lhb* mRNA in the pituitary of females are not consistent with the decrease in *lhb*-receptor mRNA observed in ovaries (Rajeswari and Unniappan 2020). This may be because *lh-receptor* expression in the ovaries is down-regulated and undergoes accelerated degradation in response to the Lh surge (Menon, Nair et al. 2007; Menon and Menon 2014). Using a zebrafish specific anti-Pnx antibody, we found that Pnx expression starts during puberty, approximately 45 days post fertilization (**Supp. Fig. 2**; (Singleman and Holtzman 2014). With the confirmation of the zebrafish Pnx protein expression profile, we then used CRIPSR/Cas9 to generate a 10 bp deletion in exon 2 of the *pnx* gene, which caused a frameshift mutation with a predicted loss of the zPnx-20 function, (*pnx^uvΔ10/uvΔ10^*) (**Fig. 1.1**). Consistent with this prediction, homozygous *pnx^uvΔ10/uvΔ10^* mutants showed a loss of zPnx immunoreactivity in the hypothalamus and pituitary when compared to non-mutants (**Fig. 1.2, A-D**). Analysis of fertility and fecundity of the heterozygous *pnx^+/uvΔ10^* and of homozygous *pnx^uvΔ10/uvΔ10^* animals did not reveal defects in reproduction (**SUPP Fig. 3**), although to date all adult fish homozygous for the *pnx* mutation have been male (**Fig. 1.3**), suggesting that *pnx* may play a role in the process of sex-determination. Further analysis of the gonads of male and female *pnx^+/uvΔ10^* heterozygous animals and of homozygous *pnx^uvΔ10/uvΔ10^* mutant males did not identify any gross morphological defects (**Supp. Fig. 4**). Thus, the loss of *pnx* function does not affect fertility.

**Figure 1:**
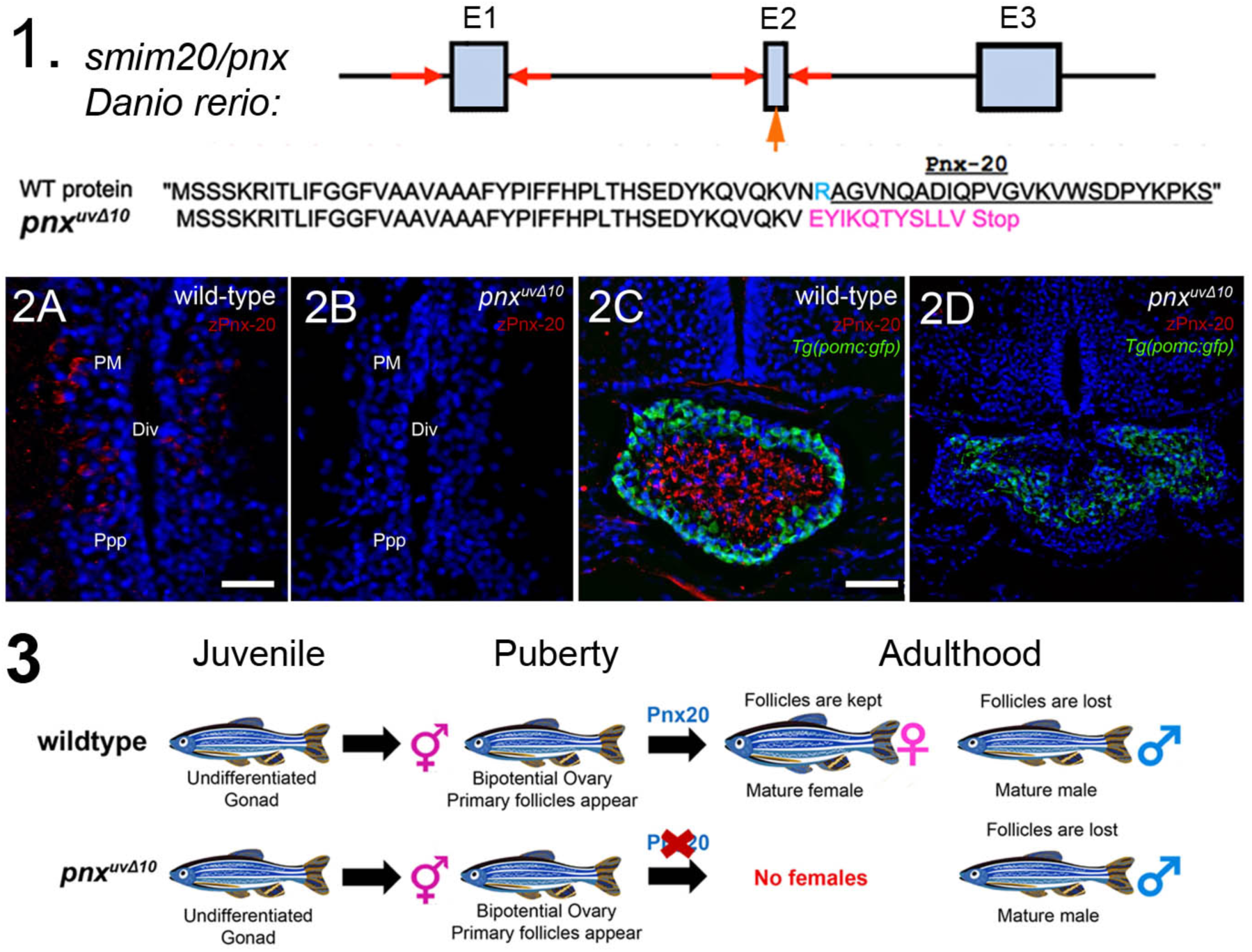
Loss of Pnx-20 function does not affect male fertility but prevents the production of female zebrafish. **1**) Location of guide RNAs and primers used to screen for mutations in Pnx-20 (red arrows); boxes represent exons (E1-3), with amino acid sequences of the wild-type *pnx* and of *pnx^uvΔ10/uvΔ10.^* protein products. **2**) Pnx-20 immunolabeling in the hypothalamus (**2A, red**) and pituitary (**2C, red**) of wildtype animals. No immunolabeling was observed in the hypothalamus (**2B**) or the pituitary (**2D**) of *pnx^uv10/uvΔ10^* homozygous mutant fish (n=3). All images are from cryostat-sectioned tissue, imaged with a z-stack of 9µm. Scale-bar= 25 µm (A, C). Magnocellular preoptic nucleus (PM), parvocellular preoptic nucleus (Ppp), adjacent to the diencephalic ventricle (DiV). **3**) Until puberty (45 dpf), a bipotential ovary can produce males or remain as female. No adult *pnx^uv10/uvΔ10^* homozygous mutant females were observed.

### Do zebrafish regulate reproduction through exogenous Gnrh?

Next we returned to the Gnrh signaling pathway and approached it from the perspective of the natural world, where hormones released into the water can act as pheromones and play an important role in fish reproduction (Sorensen, Hara et al. 1991). For example, in goldfish, 17α,20β-dihydroxy-4-pregnen-3-one (17,20β-DHP), an oocyte maturation-inducing hormone, was found to act as a pheromone. Indeed, when 17,20β-DHP is released into the water by ovulating females it triggers milt production and the initiation of sexual behavior in males (Stacey, Sorensen et al. 1989). In addition, females synthesize prostaglandin F2a (PGF_2α_) during ovulation, which can also act as a pheromone (Sorensen, Hara et al. 1988) and induce behaviors leading to successful oviposition (Kobayashi and Stacey 1993). The PGF_2α_-_’_induced response is mediated by olfactory sensory neurons (OSNs) (Sorensen and Sato 2005) and can activate specific OSNs expressing the prostaglandin receptor, which relays information via the olfactory bulb to the ventral telencephalon and the hypothalamic preoptic area (Yabuki, Koide et al. 2016); for reviews see (Stacey, Chojnacki et al. 2003; Stacey and Sorensen 2009; Stacey 2011). Similarly, in zebrafish, Chen & Martinich (1976) and Lambert *et al*. (1986) observed that females ovulate in the presence of males and that both the ovulation and the mating are controlled by pheromones synthesized by the gonads (Chen and Martinich 1976, Lambert, van den Hurk et al. 1986). In addition, ovulation could be induced by testicular homogenates, and fish rendered anosmic by sealing their nostrils did not ovulate when transferred to water that had held male fish (Chen and Martinich 1976, van den Hurk, Schoonen et al. 1987). Finally, it has also been shown that zebrafish use hormones as pheromones to initiate their sexual behavior (Lambert, van den Hurk et al. 1986, van den Hurk, Schoonen et al. 1987). However, the identity of these pheromones is unknown.

Since zebrafish are schooling fishes, we investigated whether Gnrh released into the water by conspecifics could activate the reproduction pathway in this species. To avoid any stimulation from hormones previously released by other fish into our recirculating fish housing system, the adult animals were housed individually, isolated from the housing system, and maintained in artificial fish water for 10 days. We then challenged them by adding synthetic Gnrh3 to the water (final concentration 10^-9^ M), sacrificed them after 30 minutes, and measured the levels of pituitary gonadotropins by qRTPCR. As shown in **Figure 2**, exogenous Gnrh3 increased the expression of *fshb* (**Fig. 2A**) and *lhb* (**Fig. 2B**) mRNAs in female zebrafish but not in males (**Fig. 2C** and **Fig. 2D**, respectively). These results suggest a role for exogenous Gnrh in the control of reproduction in female zebrafish. Since both the androgen receptors (Gorelick, Watson et al. 2008), and the Gnrh-Rs (Whitlock, Illing et al. 2006; Corchuelo, Martinez et al. 2017), are expressed in the olfactory epithelia (in addition to being widely expressed in other neuronal as well as non-neuronal tissues), our results support the hypothesis that exogenous Gnrh3 may activate reproduction via the olfactory organ. It has previously been shown that female zebrafish made anosmic by thermocauterizing the nasal epithelium do not ovulate when presented with water that has housed males (van den Hurk, Schoonen et al. 1987) suggesting a role for olfaction in the control of reproduction in zebrafish. To determine whether Gnrh3 signaling is mediated by the olfactory epithelia we used the protocol of (Mathuru, Kibat et al. 2012) in order to block the nares (openings) of the nose. After a 24-hour recovery period, the fish were challenged with exogenous Gnrh3, sacrificed and their pituitaries removed. Unlike the increases seen in intact females, females with blocked nares did not show increases in *fshb* (**Fig. 2A1**) or *lhb* (**Fig. 2B1**) expression when challenged with exogenous Gnrh3 (in fact these levels decreased relative to those of the controls). As a negative control zPnx-20 was added to the water at a concentration of 100 nM, a concentration that is known to elicit cellular responses (Yosten, Lyu et al. 2013; Rajeswari and Unniappan 2020) and no responses were elicited (**Supp. Fig. 5**).

**Figure 2:**
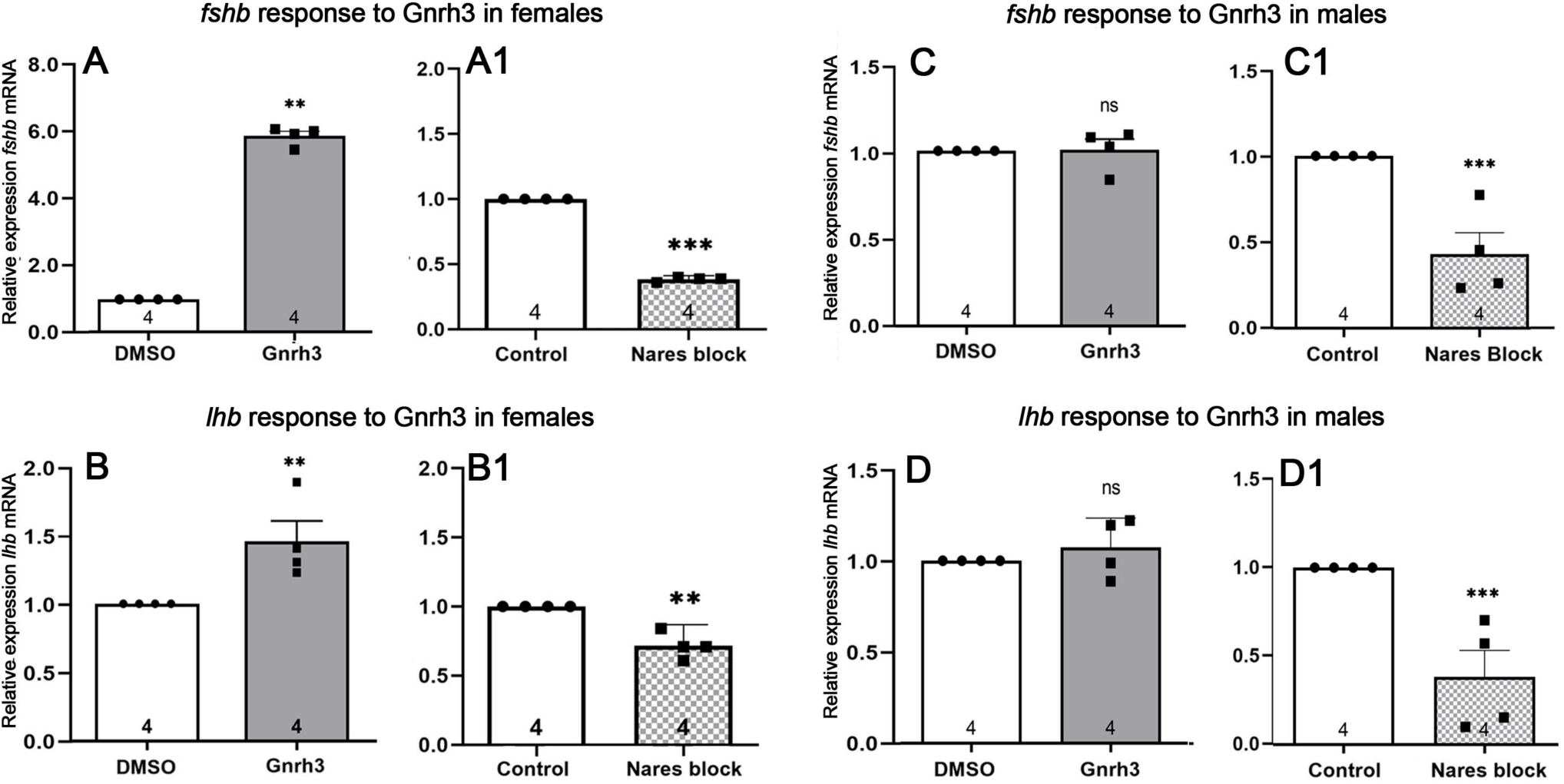
Addition of Gnrh3 to the water activates the hypothalamic pituitary axis of zebrafish. Exogenous Gnrh3 increased expression of *fshb* (**A**) and *lhb* (**B**) mRNA in wild-type female zebrafish. Blocking the nares (**A1**, **B1**) reduced the induced response. Exogenous Gnrh3 did not affect mRNA expression of *fshb* (**C**) or *lhb* (**D**) in wild-type males, but blocking the nares (**C1**, **D1**) significantly decreased the expression of gonadotropins when compared to controls. Results are expressed as mean *±* SEM and compared with the control group using *One-sample t-test*. Housekeeping gene was *beta-actin*. *\*P< 0.05; **P<0.002; ***, P<0.0002*. N=4 samples (where each sample represents pooled hypothalamus and pituitaries from 3 fish). Nares block: females N=4, males N=4.

Males, which showed no significant responses to exogenous Gnrh3, showed decreases in both *fshb* (**Fig. 2C1**) and *lhb* (**Fig. 2D1**) expression relative to controls when their nares were blocked. These results indicate that in males exogenous Gnrh3 may trigger the testosterone-based negative feedback loop on the hypothalamic pituitary axis, thereby shutting down release until the Gnrh peptide is broken down (Zhai, Shu et al. 2018). Alternatively the drop in Gnrh could be a result of the reported effects of 17α,20β-dihydroxy-4-pregnen-3-one (DHP) in the neuroendocrine regulation of reproduction in male zebrafish, where DHP is correlated with a direct stimulatory effect on pituitary *fshb* mRNA levels (Wang, Liu et al. 2016).

Thus, blocking only the nares of females and males eliminated the hypothalamic responses to exogenous Gnrh3, albeit potentially by different mechanisms in the two sexes, supporting the hypothesis that the olfactory organ mediates this sex-specific response. These results are in agreement with the original findings of Chen and Martinich (1976) where they initially observed that natural spawning of zebrafish could be induced by the introduction of ripe (ready to spawn) individuals of both sexes into a container with dechlorinated tap water, thus suggesting the presence of a waterborne trigger (Chen and Martinich 1976). In fact, these authors go as far as suggesting that Gnrh could be the relevant hormonal signal. Interestingly, (Chen and Martinich 1976) also observed that light alone (a stimulus that most zebrafish researchers consider to be the dominant trigger for spawning) does not induce the spawning of females held in isolation.

In goldfish, it has been shown that exogenous (mammalian) GnrH can be absorbed from the intraperitoneal cavity (provided by injection), from the surface of the gills (provided by careful application), and importantly for the results presented here, from the surrounding water (through bath application). In the latter case, exogenous (mammalian) GnRH was absorbed from the surrounding water and was detectable in the blood within 4 minutes of exposure. Furthermore the majority of the GnRH disappeared from the water within 1 week (50% had disappeared by Day 3), but did so more rapidly if fish were present (Sherwood and Harvey 1986).

These original observations from the literature coupled with our results may also explain why *gnrh2* and *gnrh3* loss of function mutants (Spicer, Wong et al. 2016; Marvel, Spicer et al. 2018) are fertile. Indeed, in these experiments the mutant animals were housed individually but still shared water with their conspecifics, raising the possibility that exogenous Gnrh released into the water by *wild-type* conspecifics may have rescued the sterility of the *mutant* fish.

### Loss of Gnrh-R3 gnrh-r3^uvΔ750^ reduces fecundity and fertility in zebrafish

To further explore the role of exogenous Gnrh signaling in zebrafish reproduction, we next targeted Gnrh receptor function. The zebrafish Gnrh-R1 is less sensitive to physiological doses of Gnrh peptides than is Gnrh-R3. (This is most likely due to a valine to glutamic acid substitution in extracellular loop 3, a site that is important for ligand binding in fish (Tello, Wu et al. 2008). For this reason we chose to knockout the *gnrh-r3* gene and used CRISPR/Cas9 to produce a 750 bp deletion loss of function mutation that included 5’ untranslated sequences as well as exon 1 (**Fig. 3A: Supp. Fig. 6**).

**Figure 3:**
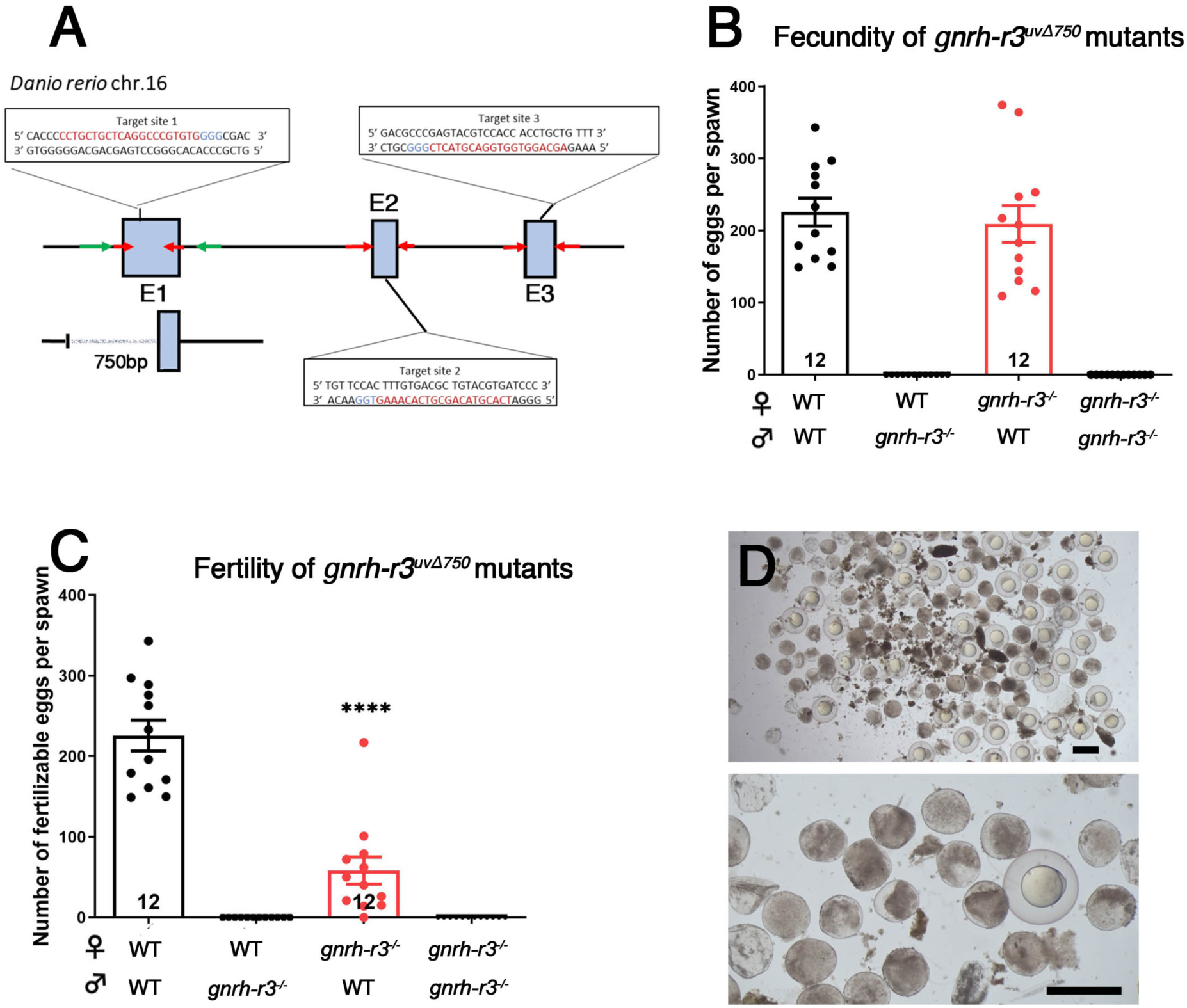
Loss of Gnrh-R3 (*gnrh-r3^uvΔ750^*) renders zebrafish infertile. **A**) Guide RNA in exon 1 of *gnrh-r3* and primers used to screen for mutants (green arrows). **B)** Crosses of wild-type females with *gnrh-r3^uvΔ750/uvΔ750^* homozygous mutant males did not produce eggs; in the reciprocal cross **(C)**, females laid eggs but most were not fertilizable. **D**) Degenerated eggs and a few fertilized embryos produced by homozygous mutant females. Scale bar= 1000µm. Data are presented as mean ± SEM; n=12 replicas per cross. Statistical significance was determined by *One-sample t-test *P< 0.0332; **P<0.0021; ***P<0.0002; ****P<0.0001*.

As shown in **Figure 3**, *gnrh-r3* mutant fish showed a striking phenotype. Indeed, *gnrh-r3^uvΔ750/uvΔ750^* homozygous mutant males were totally infertile (**Fig. 3B**) and did not elicit a mating response from wild-type females. Similarly, *gnrh-r3^uvΔ750/uvΔ750^*homozygous mutant females were mostly infertile: although they could produce eggs (**Fig. 3 B)**, most of them degenerated and were thus non-fertilizable (**Fig. 3C, D**).

Analysis of the gonads (**Fig. 4**) showed that *gnrh-r3^uvΔ750/uvΔ750^*homozygous mutant males had malformed testes that did not reach the cloaca of the animal (**Fig. 4D**, D1). In females, we found that late-stage oocytes from the ovaries of *gnrh-r3^uvΔ750/uvΔ750^* homozygous mutant female were smaller and defective (**Fig. 3B, B1**), with later stage oocytes showing degeneration (**Fig. 2D, 3B1)**. Thus, while elimination of the Gnrh3 or Gnrh2 ligands did not affect fertility or fecundity in zebrafish, the loss of function *gnrh-r3* caused almost complete sterility. These data suggest that the control of reproduction can be accomplished via exogenous Gnrh3 ligand acting on Gnrh-R3 present in the olfactory organ.

**Figure 4:**
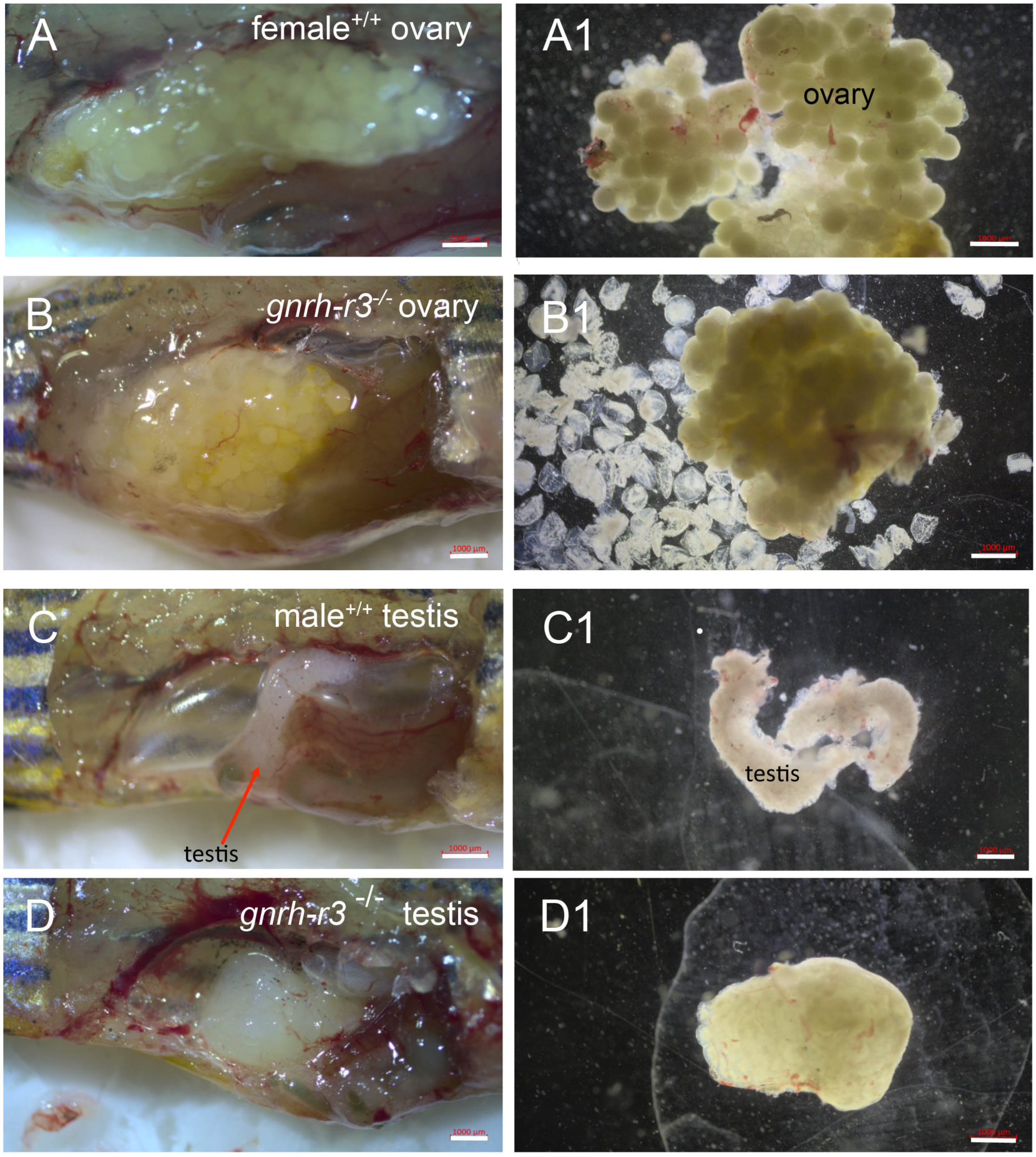
*gnrh-r3^uvΔ750^* mutants show grossly abnormal gonadal development. Gonads from intact (A-D) and dissected (**A1-D1**) wild-type fish (**A**, **C**) and from *gnrh-r3^uvΔ750/uvΔ750^* homozygous mutant animals (**B**, **D**). *gnrh-r3^uvΔ750/uvΔ750^* homozygous mutant females had irregularly shaped gonads (B) and when dissected released degenerated oocytes (**B1**). *gnrh-r3^uvΔ750/uvΔ750^* homozygous mutant males had malformed testes (**D**) and lacked sperm (**D1**), (see also **Supp. Fig. 7**). N=3 fish; Scale bars=1000 µm.

### Gnrh3 can be detected in the water housing zebrafish

Our hypothesis that GnR3 released into the water can trigger reproduction in conspecific zebrafish requires that Gnrh3 is present in the water that houses the fish. To determine whether the water housing our research animals contained Gnrh, as suggested by the original experiments of Chen and Martini (1975), we used HPLC to identify compounds present in the water. As shown in **Figure 5**, synthetic Gnrh3 had a retention time of 13.115 minutes (**Fig. 5A**, red arrow), which matched the retention time (13.114 minutes) for a prominent peak present in the circulating zebrafish water sample (**Fig. 5B**, red arrow) suggesting the presence of Gnrh3 in the circulating water. The spectrum at 280nm of the peak of the standard (**Fig. 5C**) and of the circulating fish water (**Fig. 5D**) showed high correlation between Gnrh3 and the circulating water, confirming that the compound detected in the circulating water was Gnrh3. The linear regression between peak area ratios and concentrations (0.125–10 µg/mL) (**Supp. Fig. 8**) showed excellent linearity, with a correlation coefficient (R^2^) of 0.9988. The variation coefficients for repeatability and reproducibility were 2.34% and 2.42%, respectively. The limit of detection (LOD) was 0.0625µg/mL and the limit of quantification (LOQ) was 0.2033µg/mL Gnrh3 concentration in the zebrafish system water was estimated to be 0.034µg/mL (±0.0009) corresponding to 28.05 nM (±0.72). Importantly, this concentration is within the range of concentrations of Gnrh3 needed to activate Gnrh-R3 in cell culture assays (10^-6^-10^-9^ M; (Tello, Wu et al. 2008;Cortes-Campos, Letelier et al. 2015).

**Figure 5.**
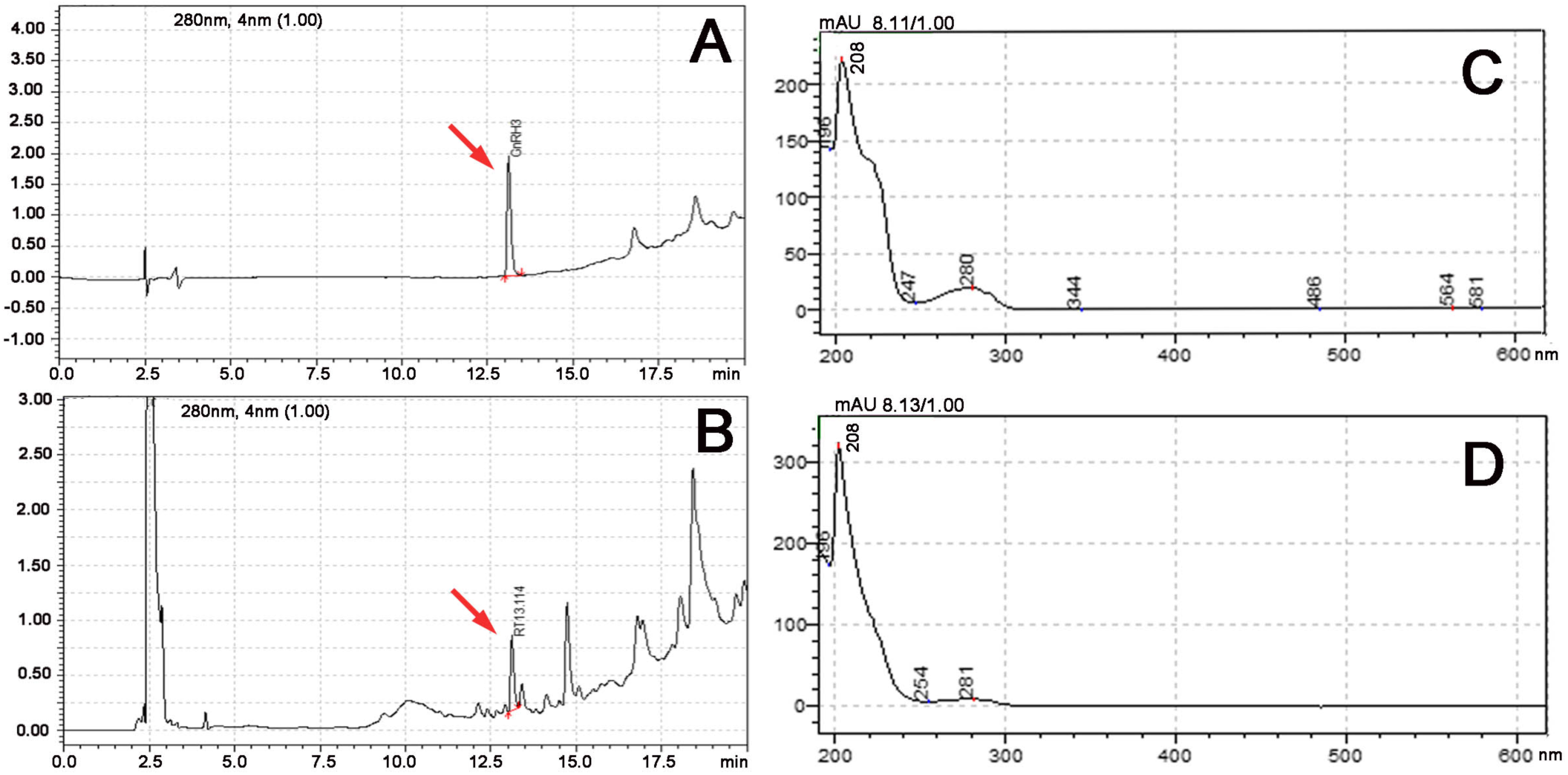
Chromatographic detection of Gnrh3 in water housing zebrafish. **A**) Synthetic Gnrh3 peptide had a retention time of 13.115 minutes (red arrow). **B**) Water sample from water housing zebrafish showed a corresponding peak at 13.114 (red arrow), matching the retention time of synthetic Gnrh3. C, D) Concordance of spectrum at 280nm for Gnrh3 peptide (**C**) and of water housing zebrafish (**D**).

This is the first report of biologically relevant levels of Gnrh in water housing zebrafish colonies, and supports the proposed hypothesis that *exogenous* GnRH plays a role in the maintenance of fertility and fecundity in the absence of *endogenous* GnRH.

## Discussion

### Gnrh signaling and the control of reproduction

The results presented here are consistent with recent reports of a dual Gnrh model suggesting that in fishes reproduction is controlled by both Gnrh and the satiety hormone Cholecystokinin (Cck). In this model, the upstream regulator of Fsh is cholecystokinin (Cck) and that of Lh is Gnrh (Uehara, Nishiike et al. 2024; Cohen, Cohen et al. 2024). In females, folliculogenesis is controlled by Fsh release, which is triggered by Cck (**Fig. 6A**, purple), and only the final process of oocyte maturation and of ovulation are controlled by Gnrh, which triggers the release of Lh from the pituitary (**Fig. 6A**, orange). In males (**Fig. 6B**), by contrast, Gnrh is the key hormone in the regulatory network that controls testes formation and spermatogenesis. In agreement with this model, successful spawning was observed in *ccka/cckb* double knockout mutant males (because testes development would be controlled by Gnrh), while no spawning was observed in *ccka/cckb* double KO mutant females (because follicular development would be under control of Cck). Yet, the *ccka/cckb* KO animals did eventually start to spawn, which the authors attribute to the existence of a compensatory mechanisms involving Lh (Gnrh upstream) (Hollander-Cohen, Cohen et al. 2024). Alternatively, our results suggest that this slow onset of spawning could be due to the presence of Gnrh in the water system. Regardless of the endocrine basis of this onset of spawning, the Cck/Gnrh pathway appears to include some degree of crosstalk, at least in zebrafish.

**Figure 6.**
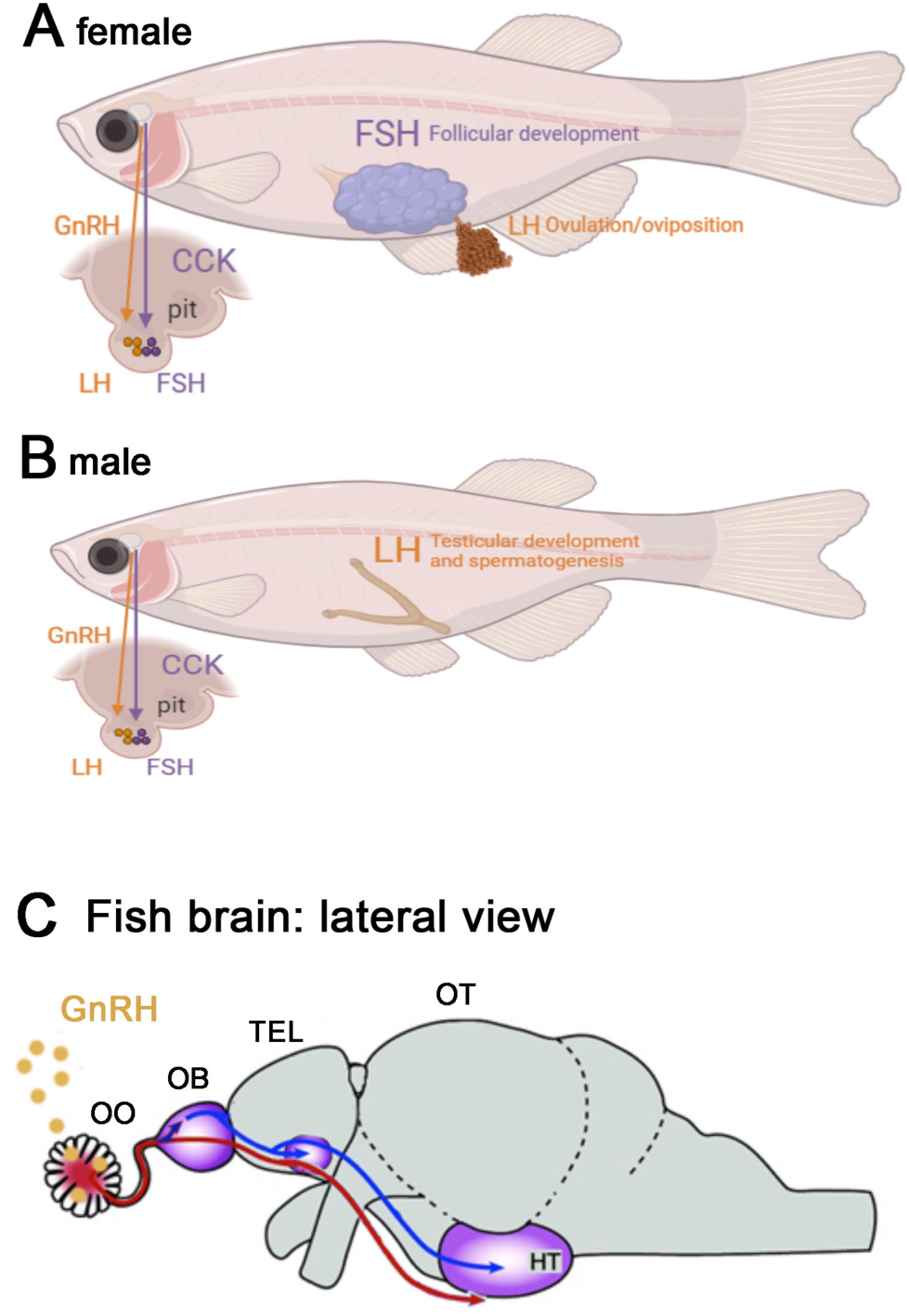
Proposed model for Gnrh-R3 control of reproduction in zebrafish. **A**) Loss of *gnrh-r3* function affects ovulation but not follicular development. In contrast in males (**B**) both testicular development and spermatogenesis are affected, consistent with current models. **C**) Exogenous Gnrh3 enters the olfactory organ (OO) where it would stimulate neuronal pathways via Gnrh-R3 (blue) or it would enter the blood through the blood vessels (red), and act on the pituitary via the blood vasculature (red tract).

In the Cck/Gnrh model, Gnrh controls the maturation of oocytes, which is consistent with the phenotype of *gnrh-r3* mutant females presented here, where the lack of Gnrh signaling results in few mature oocytes in the adult females. Our results are also consistent for the role of Gnrh in males, since animals mutant for *gnrh-r3* lack testes and do not produce sperm. Here, we have discovered the missing piece to the puzzle, namely an exogenous source of the Gnrh that can control reproduction in zebrafish.

### Potential pathways of activation

The data presented here support a role for exogenous Gnrh3 in the control of reproduction via the olfactory organ as reported by Chen and Martinich (1976) and are in agreement with the localization of Gnrh-Rs to the olfactory organ as shown by antibody staining (Whitlock, Illing et al. 2006) and mRNA expression (Corchuelo, Martinez et al. 2017). Regarding the Gnrh3 ligand signaling, the peptide could potentially act through neuronal pathways (**Fig. 6C, blue**), as described for the pheromonal PGF2α-induced reproductive responses in zebrafish (Yabuki 2016). Alternatively, because adult zebrafish have an extensive blood vasculature associated with the olfactory organ (Palominos, Calfún et al. 2022), it could pass through the blood capillaries into the vasculature that extends posteriorly to the hypothalamus (**Fig. 6C, red**). In this theoretical olfactory portal system Gnrh3 would pass into the blood capillary system via fenestrations of the olfactory organ, similar to those observed in mammals (Schaeffer, Hodson et al. 2011), and once in the blood would be rapidly transported to the pituitary (Thorne, Pronk et al. 2004). This hypothesis is supported by studies in frogs (Julien, Kouba et al. 2019) and humans (Iwamoto, Yoshida et al. 2009; reviewed in Noel and Kaiser 2011), where nasal application of Gnrh can trigger sperm production and increase serum gonadotropin, potentially through Gnrh and Fsh receptors within the vasculature (Chegini, Rong et al. 1996; Stilley and Segaloff 2018).

### Gnrh as pheromone

The term hormonal pheromone describes the situation where hormones excreted from the fish’s body act as water-borne odorants that can induce in conspecifics specific physiological and/or behavioral responses related to reproduction. (Stacey and Sorensen 2009). Here we show that the system water in which zebrafish are housed contained nanomolar concentrations of Gnrh3 peptide, raising the question of the role of waterborne hormone in the reproduction of zebrafish in the wild. Zebrafish are native to South Asia where they are found in slow-moving or still waters, such as ponds and rice fields (Engeszer, Patterson et al. 2007). In addition, zebrafish prefer shallow areas to lay large clutches of eggs, suggesting that larvae typically develop in close proximity to conspecifics (Sessa, White et al. 2008). Taken together, the studies suggest that zebrafish exist in an ecosystem where exogenous Gnrh signaling could be transmitted to conspecifics. Thus, the original studies showing that loss of function mutations in Gnrh peptides do not affect fertility (Spicer, Wong et al. 2016; Marvel, Spicer et al. 2018) may in fact be a novel example of environmental compensation for gene loss, or reveal the existence of a potential backup system in cases where endogenous Gnrh signaling fails.

### Conclusions

By considering the natural history of fishes, we uncovered a role of exogenous Gnrh in the maintenance of fertility in zebrafish. This new mechanism for Gnrh regulation of reproduction in fishes was elucidated by the surprising phenotype of the loss of function *gnrh-r3* mutant, namely profound infertility in males (extremely malformed testes) and greatly reduced fertility in females (few mature oocytes). These results are consistent with original studies suggesting that hormonal pheromones control aspects of reproduction in fishes and that exogenous Gnrh can rapidly pass into the bloodstream. The discovery that exogenous Gnrh may play an essential role in the regulation of reproduction, potentially through the blood vasculature of the olfactory organ, opens the possibility of an “olfactory-portal system”. Yet more studies are needed to determine whether there is both a blood vascular and a neural component involved in the olfactory organ-mediated response. Finally, future studies are needed determine whether the importance of exogenous Gnrh is linked to the life history of fishes (schooling versus non-schooling), and whether the ravages of climate change will impact this environmentally susceptible pathway in wild zebrafish and related species.

## Acknowledgements

We thank Trinidad Órdenes and Stephanie Alanis for the care and maintenance of the zebrafish facility at the University of Valparaiso.

## Author contributions

E. M. T. Participated in the experimental design, implementation of experiments, and writing of the manuscript; R.F., C. A-C., and I. P-B., participated in the experiments related to the reproductive system of adult zebrafish; M. E. P. participated in the experiments related to the detection of Gnrh3 in fish water. J. E. participated in analysis of genetic and genomic data and editing of the manuscript; K. E. W. Principal Investigator, participated in oversight of project, experimental design, implementation of experiments, and writing of manuscript.

## Bioethics and Animal Care Committee

Gabriela Muñoz, *Instituto de Biología, Miembro Titular, Presidenta*; Rodrigo Toro, *Instituto de Neurociencia, Miembro Titular, Vicepresidente*; María Victoria Velarde, *Instituto de Fisiología, Miembro Titular, Secretaria*; Enzo Seguel, *Dirección de Investigación, Miembro Titular, Veterinario Institucional*; Jonathan Martínez, *Instituto de Fisiología, Miembro Titular;* Leticia Luna, *Facultad* de Farmacia, *Miembro Titular*. https://sites.google.com/uv.cl/bioeticaciencias/inicio

## Data availability statement

The manuscript does not include accession codes, unique identifiers, or web links for publicly available datasets, nor are there any restrictions on data availability or clinical datasets.

## Materials and Methods

### Animals

Wild-type (WT) zebrafish of the Cornell strain (derived from Oregon AB) were used. Zebrafish were maintained in a recirculating system (Aquatic Habitats Inc., Apopka, FL) at 28°C on a light-dark cycle of 14 and 10 hours, respectively. Embryos were obtained from natural spawnings under laboratory conditions and raised at 28.5°C in Embryo medium (Westerfield 2007). Staging of pubertal fish was done according to (Singleman and Holtzman 2014). Transgenic line Tg:(*pomc:GFP*) (Liu, Huang et al. 2003) was used to visualize specific cell types. All procedures were approved by the Institutional Animal Use and Care Committee for Animal Research at the University of Valparaíso #BIO-UVFC-4.

### Cryo-sectioning

Fish of specific development stages were sacrificed, and the heads collected and fixed in 4**%** paraformaldehyde for 24 hours at 4°C. They were then decalcified in 0.2 M ethylene-diamine-tetra-acetic acid (EDTA) solution pH 7.3 for 2 days at 4°C. Tissues were embedded in 5% sucrose/1.5% agarose in MilliQ H20. Blocks were submerged in 30% sucrose overnight, embedded in O.C.T Compound (Tissue Tek®) and frozen in cryomolds at –20°C for at least 2 days prior the cryo-sectioning. Thirty micro-meter cryo-sections were processed for immunohistochemistry.

### Zebrafish anti-Pnx-20 Antibody

A polyclonal anti-zebrafish Pnx-20 antiserum was generated in rabbit by Pacific Immunology (USA) https://www.pacificimmunology.com/, against the zebrafish Pnx-20 peptide (AGVNQADIQPVGVKVWSDPYKPKS) as an immunogen. The specificity of the antiserum is demonstrated by the lack of immunostaining in the Pnx-20 mutant fish.

### Antibody Staining

Cryosections were incubated overnight at 4°C with rabbit anti-zPnx-20 specific for zPnx-20 peptide (1:500, Pacific Immunology, USA) and mouse anti-GFP (1:500, Invitrogen A-11120) primary antibodies. They were then processed using VECTASTAIN elite ABC kit (Rabbit IgG, VECTOR; PK-6101) and DAB Peroxidase (HRP) Substrate (VECTOR; SK-4105), following manufacturer instructions, or Alexa Fluor 568 conjugated anti–rabbit antibody (goat 1:500, Molecular Probes) and Dylight 488–conjugated anti–mouse antibody (goat 1:500, Jackson ImmunoResearch).

### Total RNA isolation and reverse transcription

The pituitaries and hypothalamus of 3 animals from each group were combined, and total RNA was extracted using TriZol Reagent (Ambion, Life Technologies) according to previously published protocols (Peterson and Freeman 2009). Briefly, the tissues were isolated from the adult male or female zebrafish and homogenized in 1mL TriZol-Reagent. Two hundred µL of chloroform (Sigma-Aldrich) were then added to each tube to a total volume of 1200µL and the mixture centrifuged at 4°C for 15 minutes at 12000 ×g (Sorvall legend RT). After centrifugation, the aqueous layer (around 500µL) was transferred to a new tube and the same volume of chloroform/isoamyl alcohol acidic pH was added and centrifuged at 4°C for 5 minutes at 16000 ×g. The aqueous layer was transferred in a new tube and 225 uL of NaAcetate 3M pH 7 and 800uL of ethanol 100% was added into the solution and incubated overnight at –20°C and centrifuged for 30 minutes at 16000 ×g at 4°C to precipitate the RNA (Westerfield 2007;Peterson and Freeman 2009). RNA was then washed with 80% ethanol/20% DEPC treated water. Total RNA was re-suspended with DEPC treated water and treated with DNAse I (Amplification grade, Invitrogen), then quantified by the Qubit HS RNA Assay Kit (Invitrogen, ThermoFisher Scientific) using a Qubit 3.0 Fluorometer (Invitrogen, Life Technologies). To synthetize cDNA from total RNA, reverse transcription was performed at 42°C for 1 hour in a reaction of 20µL containing x µL of total RNA at the initial concentration, 1µL of 10mM dNTP mix, 1µL of 0.5 µg of Oligo(dT) (Invitrogen), 50 U of SuperScript II reverse transcriptase (Invitrogen) and RNAse Out (Invitrogen). The cDNA was then used for qPCR.

### Quantitative PCR (qPCR) of *lhb* and *fshb* mRNAs expression

qPCR was performed using a M×3000P thermocycler (Stratagene). The reaction mixture consisted of 2µL of (cDNA), 0.75µL of each primer, and 12.5 µL of SYBR Green/ROX qPCR Master Mix (Molecular Probes), in a final volume of 25µL. The reaction profile consisted of 95°C for 10 minutes, then 40 cycles of: 95°C for 20 sec, 59°C for 20 sec, and 72°C for 20sec for signal detection. The data were analyzed using the thermocycler software (MxPro-Mx3000P v4.10) and Prism8.

All gene expression values were normalized to the housekeeping gene, *beta-actin* (Calfun, Dominguez et al. 2016). Relative expression was calculated using the 2^−ΔCt^ method relative to control levels. For statistical analyses were performed using Student *t*-test; significantly differentially expressed genes were identified as those with a *P*-value ≤ 0.05.

### CRISPR/Cas9 design

All gRNAs were designed using CHOPCHOP (https://chopchop.cbu.uib.no/) and Integrated DNA Technologies

(https://www.idtdna.com/site/order/designtool/index/CRISPR_CUSTOM). Specific sgRNAs and Constant oligonucleotide were purchased from Macrogen Inc (South Korea). sgRNAs were generated following (Gagnon, Valen et al. 2014). For each sgRNA, 60 base pair oligonucleotides were synthesized and included: the T7 promoter for in vitro transcription, the 20-base spacer region specific to the target gene, and an overlap region that anneals to the constant oligonucleotide. If the sgRNA did not start with 2 GG, these bases are added to enhance the transcription reaction of the T7 polymerase. The gene-specific oligos using T7 have the architecture TAATACGACTCACTATA-N20-GTTTTAGAGCTAGAAATAGCAAG, with Ns replaced by the 20-base specific to the target and the constant-oligo: AAAAGCACCGACTCGGTGCCACTTTTTCAAGTTGATAACGGACTAGCCTTATTTTAA CTTGCTATTTCTAGCTCTAAAAC. Annealing of sgRNA and Constant-oligo used 1uL each (100uM stock) in a total volume of 10uL under the following conditions: 5 minutes at 95°C, then 85°C reduced at –2°C per second, then 85°C to 25°C at –0.1°C/second, and a final hold at 4°C. Finally, 10 µL of solution containing 0.5 µL of T4 NEB DNA polymerase, 2.5 µL of dNTPs (10mM), 2 µL of 10X NEB Buffer2, 0.2 µL of 100X NEB BSA and 4.8 µL of water were added and incubated at 12°C for 20 minutes to fill-in. The template was purified using PCR clean-up column (Nalgene) and verified by electrophoresis. In vitro transcription of the dsDNA was run overnight using 3 µL of the purified template with T7 transcription Kit (New England Biolabs E2040S). gRNAs were then purified using RNAeasy mini Kit (Qiagen 74104).

### Primers

<u>Pnx</u>: The sgRNAs target site for *pnx* (ensemble gene ID: ENSDARG00000112670) located in exon 2 was 5’GCAAGTGCAGAAGGTGAACC. <u>Gnrh-R3</u>: The sgRNAs target site for *gnrh-r3* (GeneBank accession number: NC_007127) in exon 1 was 5’TGCTGCTCAGGCCCGTGTGG-3’.

### CRISPR/Cas9 injection and generation of mutants

The injection solution (2 nL) consisted of 50 ng/µL of sgRNA for *pnx* or 25 ng/µL sgRNAs for *gnrh-r3*, with 1 µL of Cas9 protein (TrueCut Cas9 Protein v2 5µg/µL #A36499, ThermoFisher) and 1 µL of phenol-red. The solution was heated at 37°C for 5 minutes prior to the injection to allow the formation of sgRNA-Cas9 complex and injected into the yolk of 1-2 cells stage embryos.

Founders were identified by PCR and outcrossed to wild-type fish (F1), then in-crossed to generate heterozygote and homozygote F2s. For each target site, specific primers were designed to identify carriers.

*Pnx* mutant lines were genotyped using the following primers:

Forward: 5’ CAACTGGGCCATTAGAAATCA

Reverse: 5’TACCAACAGGCTGTATGTCTGC (for exon 2_.

*Gnrh-r3* mutant lines were genotyped using the following primers:

Forward: 5’-GTGAAACTGGATCTCTCTGTCCTT-3’

Reverse: 5’-GGTGTCTGTCCAGACTGATG –3’

### Gonadal morphology

Adult zebrafish (10-12 months) were sacrificed using 400 mg/100ml tricaine (#A5040 Sigma-Aldrich)(Wilson, Bunte et al. 2009) and gonads collected and fixed in Bouin’s solution for 48 h at 4°C. They were then rinsed in 70% ethanol and dehydrated in an ethanol series, cleared in Histoclear (Sigma-Aldrich) and embedded in Paraplast Plus (Sigma Chemical Co., St. Louis, MO, USA). Serial sections (5 µm; Leica RM 2155 microtome) of the gonads were mounted on slides, de-paraffinized and rehydrated. Sections processed for histology were stained with Hematoxylin of Harris (Sigma Chemical Co, USA) and Chromotrope 2R (Sigma Chemical Co, USA). The sections were then dehydrated and mounted with Entellan (107961-Merck Millipore).

### Exogenous Gnrh3 exposure

Ten days before exposing the zebrafish to exogenous Gnrh3, the animals (n=3 per group; age and weight-matched) were housed individually in 2L tanks of artificial water that was matched for conductivity and pH with fish system water. Importantly, this water was not shared with the water in the circulating system of the fish facility. The water was replaced every day with clean water and the fish were fed twice a day with dry food. On the day of Gnrh3 exposure, the fish were first acclimatized for 1 hour in 450mL water in a new tank to minimize the background activation of the olfactory epithelium (Yabuki, Koide et al. 2016). After this acclimatization, 10^−9^M of synthetic Gnrh3 (Sigma-Aldrich #L4897) dissolved in dimethylsulfoxide (DMSO) or DMSO alone (control; Sigma-Aldrich) was added to the water. Because it has previously been shown that hormonal pheromones activate neurons between 10-30 minutes after exposure (as assayed by Immediate Early Gene expression; (Yabuki, Koide et al. 2016) the fish were sacrificed after 30 minutes, and the pituitaries collected and stored at –80°C in TRIzol Reagent (Ambion, Life Technologies) until further analysis. To avoid interaction with potential endogenous peaks of Gnrh (which occur at 14:00 to 16:00 hrs), or the subsequent transcription of *lhb* and *fshb* mRNAs (16:00 to 18:00) as reported in Medaka (Karigo, Kanda et al. 2012), hormone exposures were done in the morning between 09:00 and 11:00.

### Blocking of nares

One day before Gnrh3 exposure, adult fish were anesthetized in Tricaine (400mg/100ml; A5040 Sigma-Aldrich) and immobilized with a wet sponge in a Petri dish. The nares were then dried with Kim-wipes under a dissecting microscope and a small drop of Leukosan® Adhesive (Chemence Medical Inc, USA) tissue adhesive was applied to seal both the anterior and the posterior nares of both noses (Mathuru et al., 2012). Thirty seconds after the application of the adhesive, fish were allowed to recover overnight in their individual tanks and exposed to Gnrh3 peptide as described above.

### HPLC

The elements present in the water from our circulating water system were identified by high-performance liquid chromatography (HPLC). For this, one liter of circulating fish water was collected during the afternoon [between 15:30 to 13:45]. Two fractions of 300mL and one fraction of 200mL of the fish water were frozen at –80°C overnight. The frozen 800 mL of water were then lyophilized using a freeze dryer (INOFD-18PU, Innova), and reconstituted in 3mL or 2mL of 50:50 methanol (Merck, Germany) in UtraPure Distilled water (Invitrogen). The solution was then filtered using a 13 mm, 0.22um nylon syringe filter (SCEQ-CF2101-A, Everest Scientific). Synthetic Gnrh3 peptide (≥ 97% HPLC; L4897, Sigma-Aldrich) at a concentration of 10ug/mL was used as the standard.

Chromatographic separations were carried out using a Shimadzu HPCL system with a reverse phase analytical column Inertsil ODS-3 (150 mm x 4.6 mm x 5 um; GL Sciences Inc, Japan) and UV-VIS detector/Photodiode Array Detector (SPD-M20A, Shimadzu) at 280 nm, as described in (Gaikwad, Biju et al. 2005) with some modifications. The mobile phase contained 0.1% trifluoroacetic (TFA) in acetonitrile (eluent A) and 0.1% TFA in UltraPure H_2_0 (eluent B) at a flow rate of 0.7 mL/min. The gradient program was as follow: (time, min/%B) 0/2%, 15/98%, 20/2%, at 25°C and the injection volume was 20 µL. The standard and the samples were injected at least 3 times ea. Each injection was preceded by a blank run to ensure that the column was not contaminated. For precision purposes one sample was analyzed one day six times on one day and another six times the next day. To determine the concentration of Gnrh3 in the fish water we used a calibration curve using synthetic Gnrh3 peptide solutions in the range of 0.125–10 µg/mL. The sample and instrumental methods were validated based on *USP 47–NF 42* (2024) for their linearity, precision (repeatability and reproducibility), limits of detection (LOD) and quantification (LOQ) parameters.

## SUPPLEMENTAL DATA

**Supplemental Figure 1:**
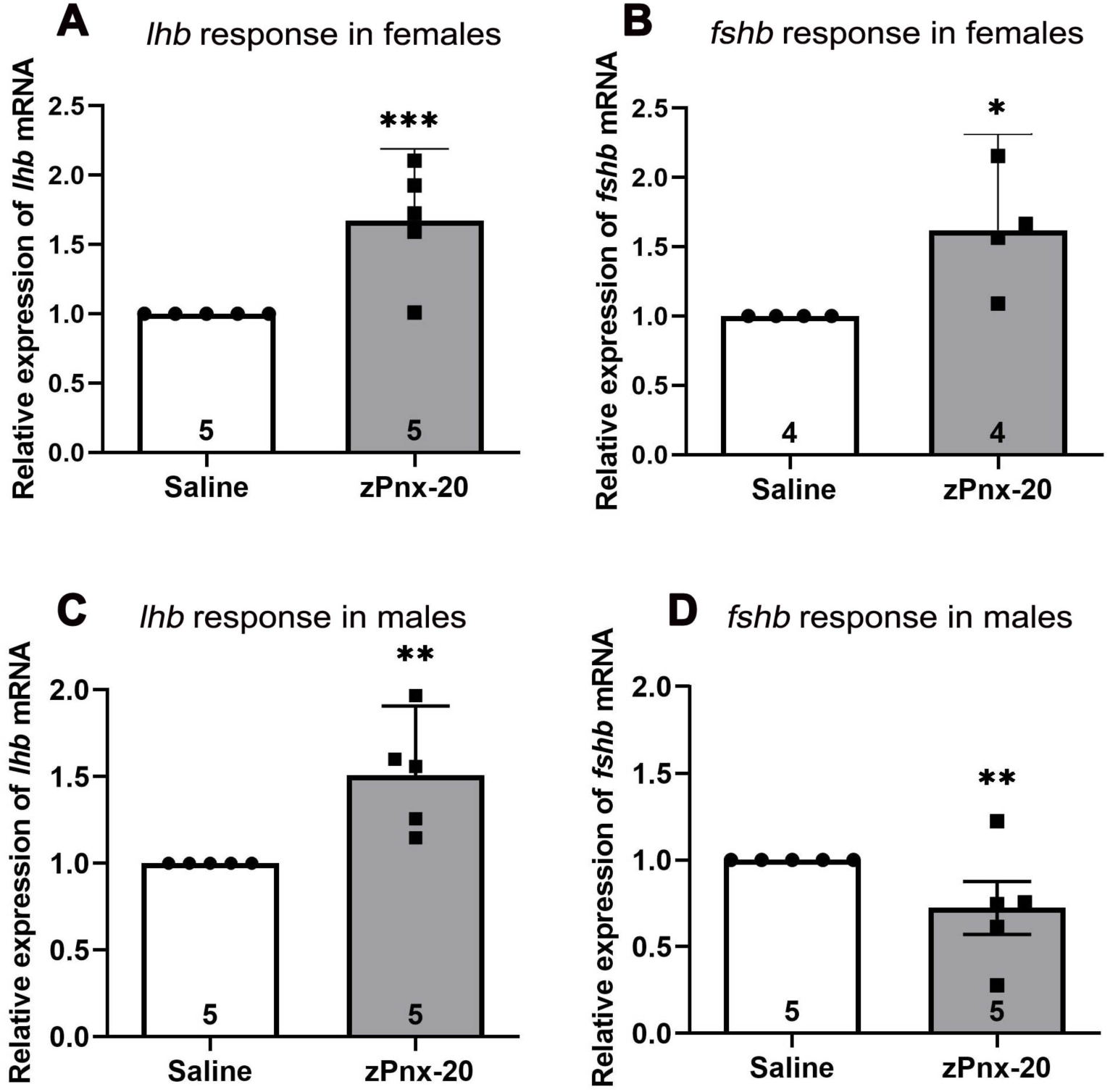
Pituitary response to zPnx-20. Relative mRNA expression of gonadotropins in response to intraperitoneal injections of zPnx-20 peptide. In adult females *lhb* mRNA (**A**) and *fshb* mRNA (**B**) were significantly upregulated in response to the zPnx-20 peptide injection. In males *lhb* mRNA (**C**) showed a significant increase in expression whereas *fshb* mRNA was downregulated (**D**). Housekeeping gene was *beta-actin*. Data: are expressed as mean ± SEM. Results from the experimental group were compared with those of the control group using a *One-sample t-test*. *\*P< 0.05, **P<0.002, ***P<0.0002*. N=5. For females, *fshb* N=4 samples (where each sample represents pooled hypothalamus and pituitaries from 3 fish).

**Supplemental Figure 2:**
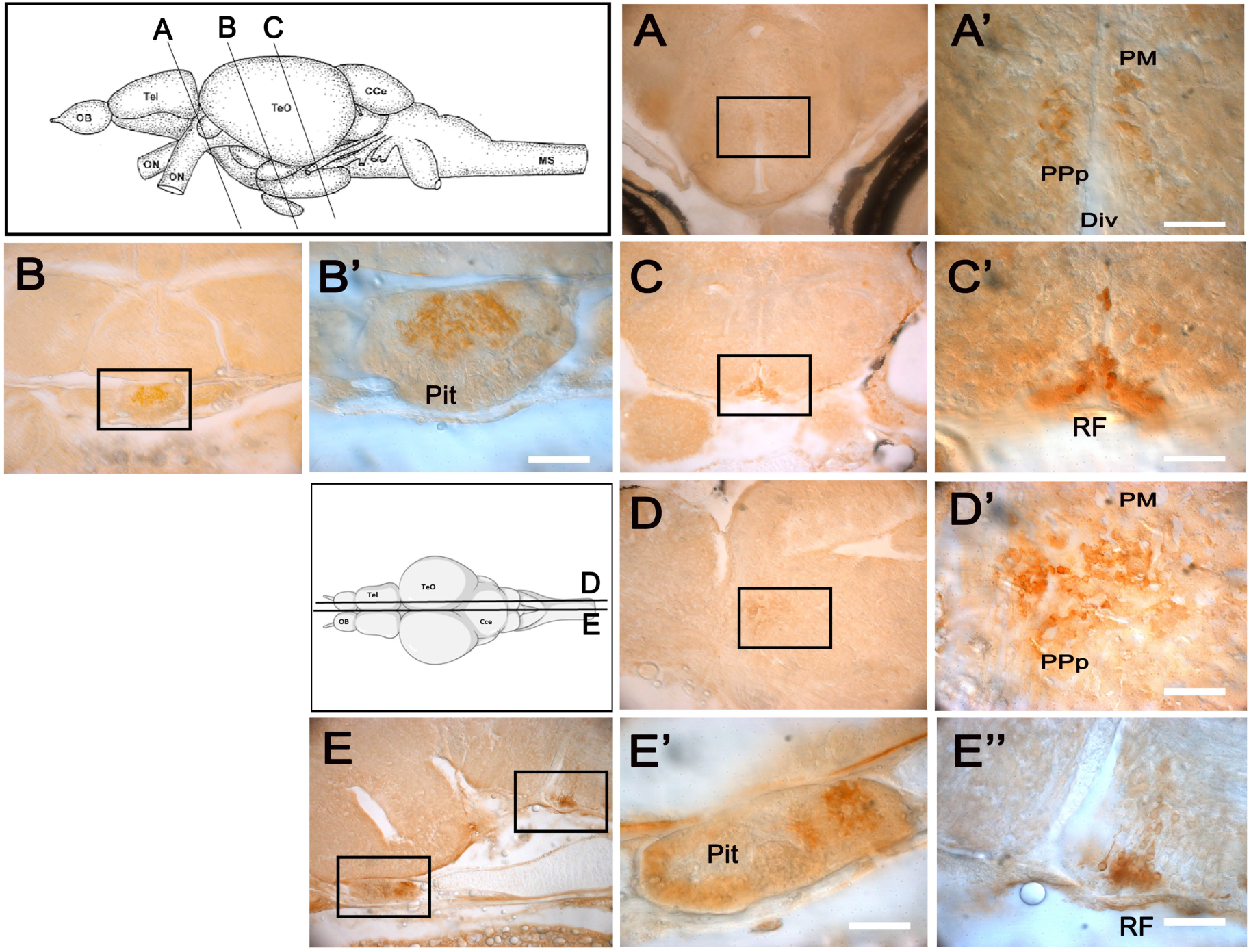
Expression of zPnx-20 in the zebrafish hypothalamus at the onset of puberty. (**A-C**) Coronal cryostat section of 47dpf zebrafish showing zPnx-20 immunoreactivity in the magnocellular pre-optic nucleus (PM) and the parvocellular pre-optic nucleus (PPp) adjacent to the diencephalic ventricle (DiV) (**A–A’**) in the pituitary (Pit) (**B–B’**) and in the Raphe (**C-C’**). A-C’ are frontal sections. (**D-D’**): Sagittal cryostat section of 47dpf zebrafish showing zPnx-20 immunoreactivity in the magnocellular pre-optic nucleus (PM) and the parvocellular pre-optic nucleus (PPp) adjacent to the diencephalic ventricle (DiV). **E-E’’**) zPnx-20 immunoreactivity in the pituitary and the Raphe (RF). **E’**) Higher magnification of boxed area in **E** (pit) and (**E’’**) higher magnification of boxed area in **E** (RF). **D-E’’** are sagittal sections. Data N=8 fish, 30uM sections; Scale-bar 50µM (**A’, B’, C’, D’, E’** and **E’’**)

**Supplemental Figure 3:**
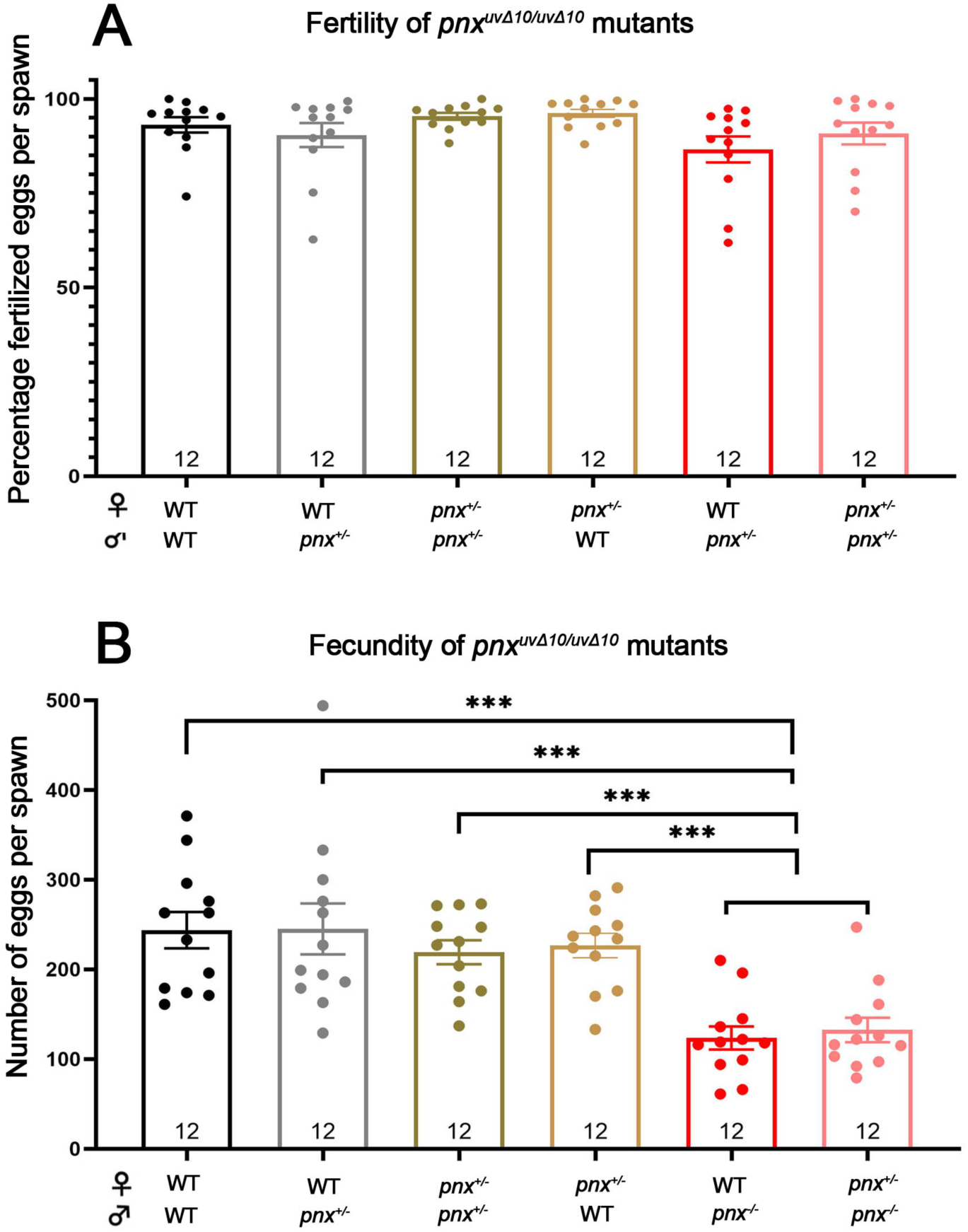
Reproductive capacities of wild-type fish compared to that of *pnx^uvΔ10/uvΔ10^* homozygous mutant fish. **A**) Number of eggs laid per cross of one male paired with one female, in which heterozygous *pnx^+/uvΔ10^*or wild-type females were crossed to homozygous *pnx^uvΔ10/uvΔ10^*mutant, heterozygous *pnx^+/uvΔ10^*, or wild-type males (no homozygous *pnx^uvΔ10/uvΔ10^* females were produced). **B**) Percentage of eggs fertilized from the crosses shown in (**A**). Data are presented as mean ± SEM; N=12 replicas per cross. Statistical significance *P-values* were calculated with *One-sample t-test* and Ordinary one-way ANOVA. *\*P< 0.05; **P<0.002 ***; P<0.001*.

**Supplemental Figure 4:**
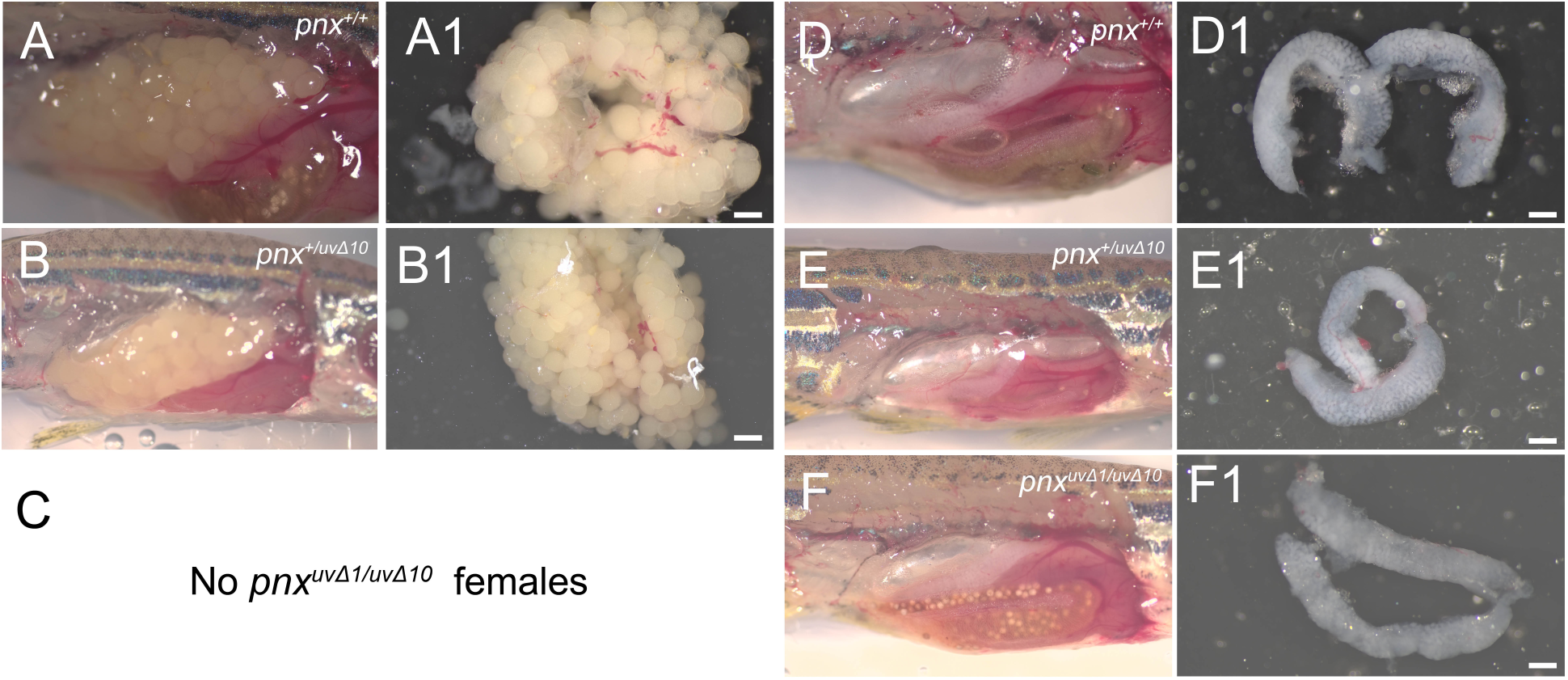
Morphology of gonads from wild-type and *pnx* mutant fish. (**A, A1**): ovaries from wild-type females and from *pnx^+/uvΔ10^* heterozygous mutant females (**B, B1**) showing normal ovary and oocyte structure. **C**) Homozygous *pnx^uvΔ1/uvΔ10^* mutant females were never observed. **D, D1**) Wild-type male testis showing normal structure and extension. **E, E1**) Testis of heterozygous *pnx^+/uvΔ10^* and of homozygous *pnx^uvΔ10/uvΔ10m^* mutant males (**F, F1**) showing normal structure. N=3 fish; Scale bars 500uM.

**Supplemental Figure 5:**
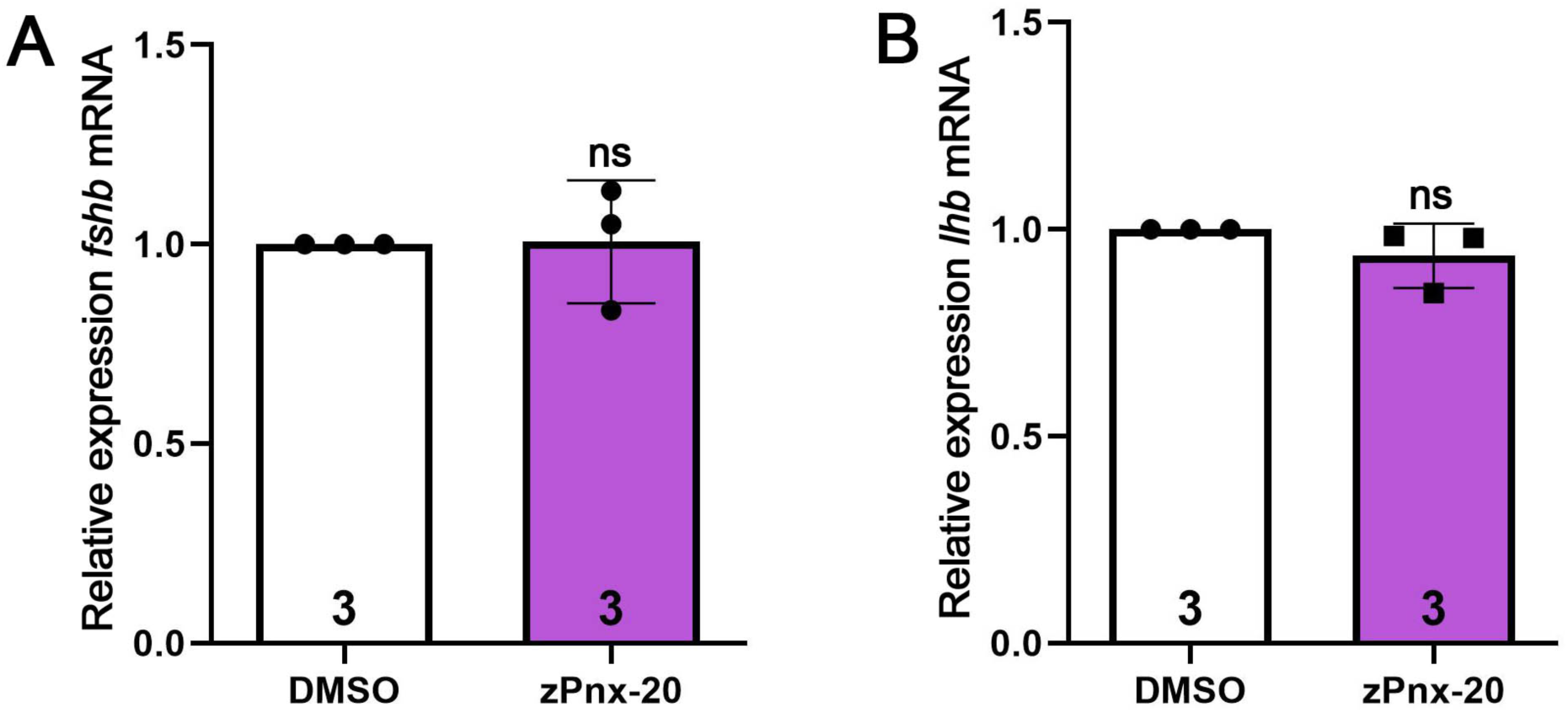
Adding 100nM Phoenixin-20 (zPnx-20) peptide to the water did not stimulate mRNA expression of pituitary gonadotropins, *fshb*. (**A**) (p=0.9467) or *lhb* (**B**) (p=0.2317), in females. Housekeeping gene was *beta-actin*. Data: are expressed as mean ± SEM. Results from the experimental group were compared with those of the control group using a *One-sample t-test*. *\*P< 0.05; **P<0.002; ***P<0.0002*. N=3.

**Supplemental Figure 6.**
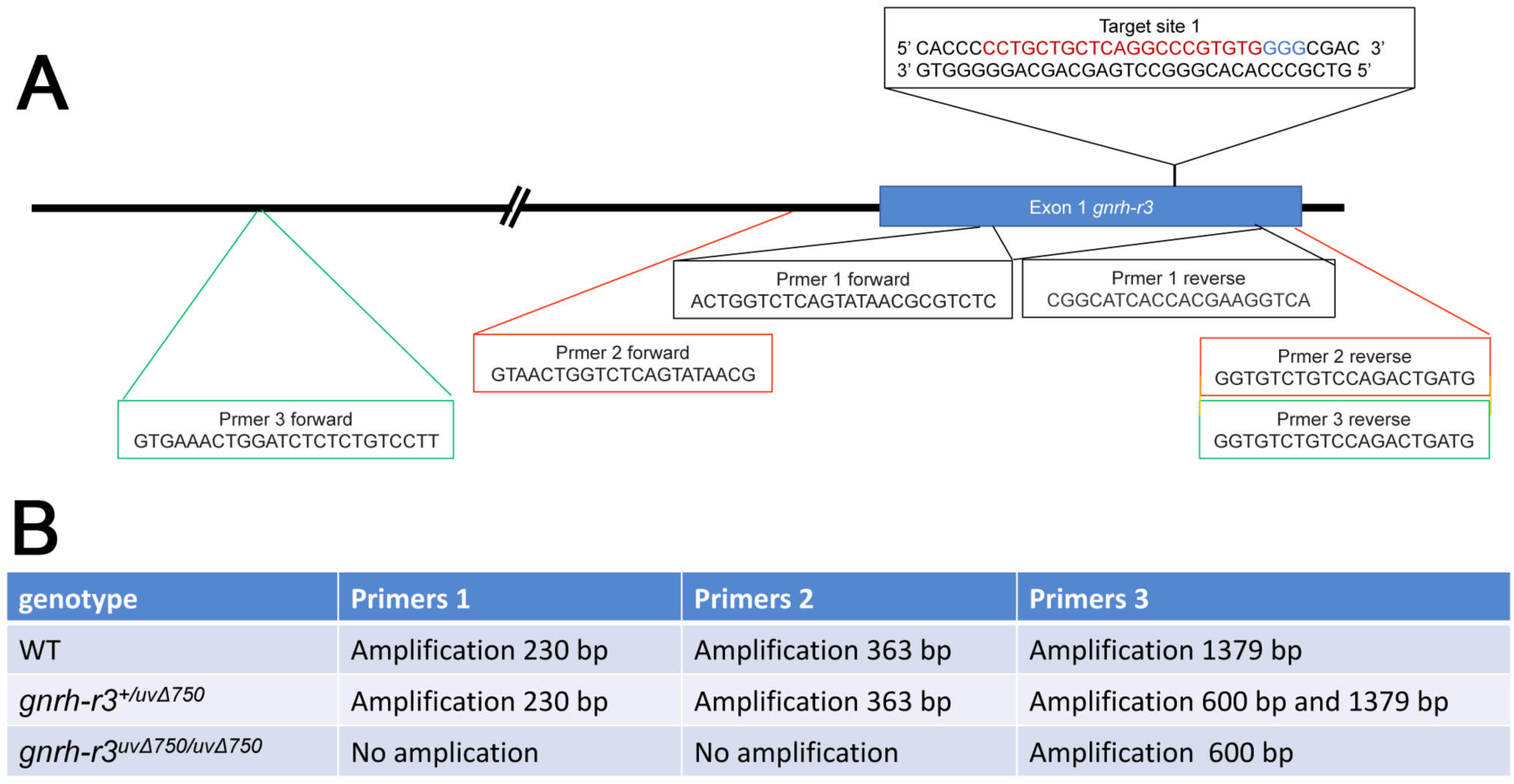
Primer Mapping of *gnrh-r3^uvΔ750/uvΔ750^* mutation. **A**) Location in WT genome (chromosome 16) of primer pairs used to map *gnrh-r3* deletion. **B**) Table showing amplicons for primer pair 1 (black), pair 2 (red) and pair 3 (green) in wild-type (WT), and mutant *gnrh-r3^+/uvΔ750^* and *pnx^uvΔ10/uvΔ750^* animals.

**Supplemental Figure 7:**
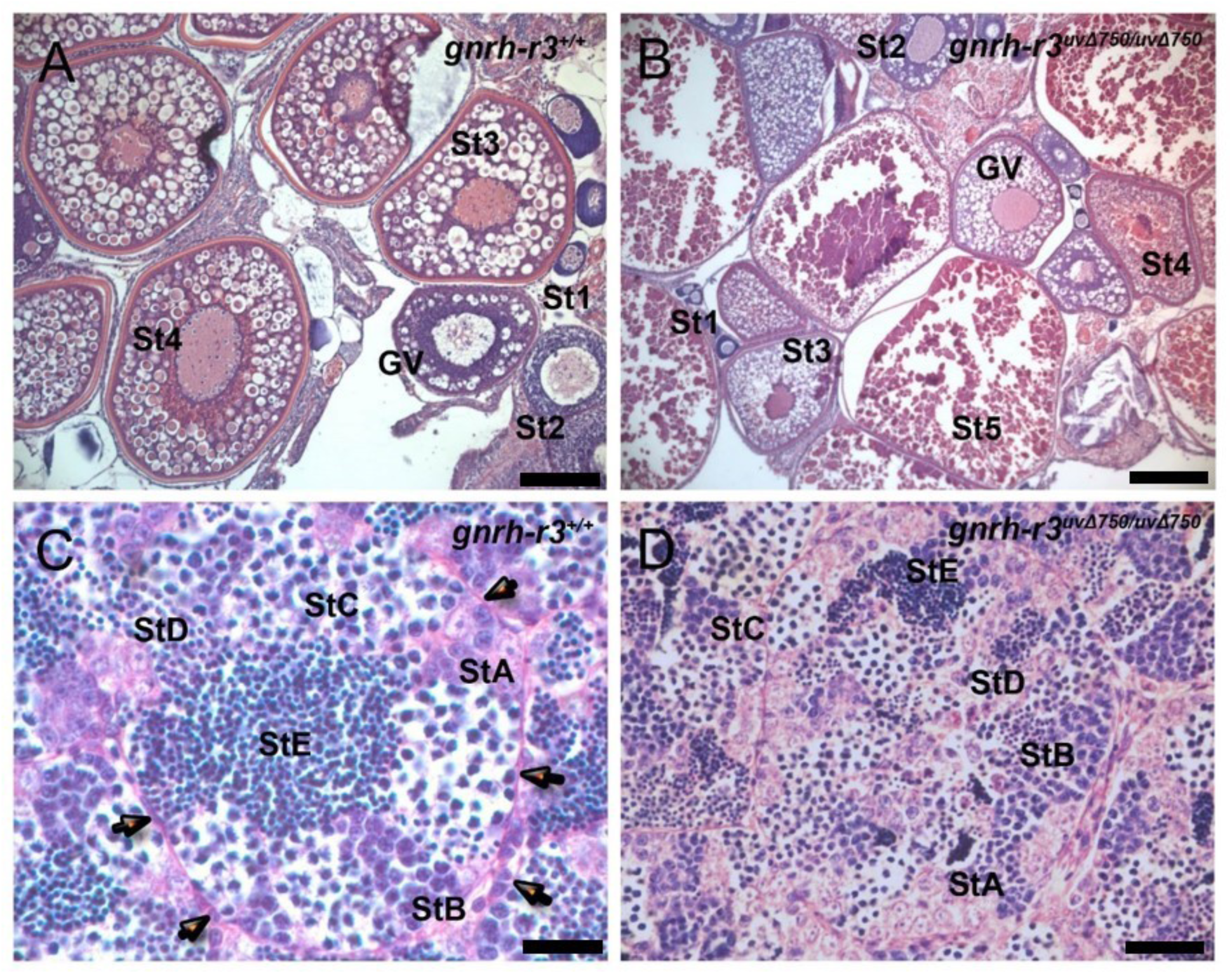
Gonadal phenotype of *gnrh-r3^uvΔ750/uvΔ750^* mutants. **A**) Ovary of wild-type female showing all the stages of oocyte development from the early stages: oogonia (St1) and early follicles stages (St2) (recognizable because of the presence of a visible germinal vesicle; GV), to the transitional stage or cortical alveolus stages (St3). **B**) Ovary of *gnrh-r3^uvΔ750/uvΔ750^* homozygous mutant female, where all the stages of oocyte development could be found but later stages (St4 and St5) showed signs of oocyte degeneration. **C**) Testes of wild-type males showing normal organization within seminiferous tubes (arrows). The spermatogonia (StA), characterized by large nuclei, are organized in clusters located in the periphery of the seminiferous tubes: Associated with the spermatogonia are the leptotene stage cells (StB), with smaller nuclei with the primary spermatocytes (StC) and secondary spermatocytes (StD) having the smaller nuclei. The spermatozoa (StE) are evident as tightly clustered very small nuclei located in the middle of the seminiferous tubes. **D**) Testis of *gnrh-r3^uvΔ750/uvΔ750^* homozygous mutant male showed abnormal seminiferous tubes, with borders difficult to define. The tissue lacked cellular organization at the different stages of maturation with the spermatogonia located throughout the testes. StA: spermatogonia, StB: leptotene stage, StC: zygotene stage primary spermatocytes, StD: secondary spermatocytes, StE: spermatozoa. N=3 fish, scale bar 30uM.

**Supplemental Figure 8:**
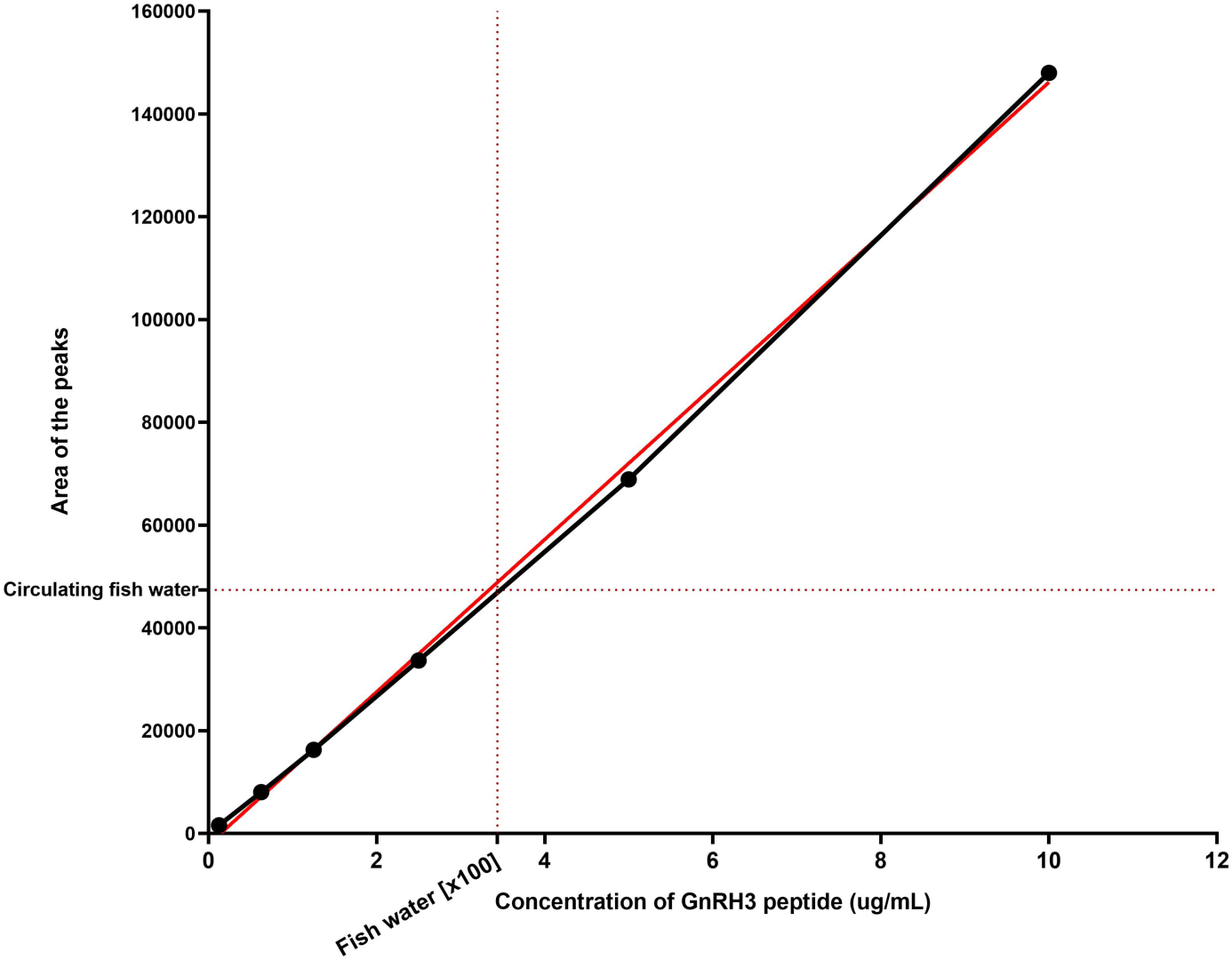
Calibration curve (black) showing the peak area (Y axis) of standard Gnrh3 vs. concentration (X axis). The red line shows the regression line calculated from the mean area for each concentration. The red dotted line on the Y axis marks the reading for the 100x concentrated Gnrh3 peptide in the circulating fish water, and was used to estimate the peptide concentration (Quattrocchi, Abelaira et al. 1992).

